# SARS-CoV-2 protein Nsp1 alters actomyosin cytoskeleton and phenocopies arrhythmogenic cardiomyopathy-related PKP2 mutant

**DOI:** 10.1101/2020.09.14.296178

**Authors:** Cristina Márquez-López, Marta Roche-Molina, Nieves García-Quintáns, Silvia Sacristán, David Siniscalco, Andrés González-Guerra, Emilio Camafeita, Mariya Lytvyn, María I. Guillen, David Sanz-Rosa, Daniel Martín-Pérez, Cristina Sánchez-Ramos, Ricardo García, Juan A. Bernal

## Abstract

Mutations in desmosomal *Plakophilin-2 (PKP2)* are the most prevalent drivers of arrhythmogenic cardiomyopathy (ACM) and a common cause of sudden cardiac death in young athletes. However, partner proteins that elucidate PKP2 cellular mechanism to understand cardiac dysfunction in ACM are mostly unknown. Here we identify the actin-based motor proteins Myh9 and Myh10 as key PKP2 interactors, and demonstrate that the expression of the ACM-related PKP2 mutant R735X alters actin fiber organization and cell mechanical stiffness. We also show that SARS-CoV-2 Nsp1 protein acts similarly to this known pathogenic R735X mutant, altering the actomyosin component distribution on cardiac cells. Our data reveal that the viral Nsp1 hijacks PKP2 into the cytoplasm and mimics the effect of delocalized R735X mutant. These results demonstrate that cytoplasmic PKP2, wildtype or mutant, induces the collapse of the actomyosin network, since shRNA-*PKP2* knockdown maintains the cell structure, validating a critical role of PKP2 localization in the regulation of actomyosin architecture. The fact that Nsp1 and PKP2 mutant R735X share similar phenotypes also suggests that direct SARS-CoV-2 heart infection could induce a transient ACM-like disease in COVID-19 patients, which may contribute to right ventricle dysfunction, observed in patients with poor survival prognosis.

**Highlights:** The specific cardiac isoform Plakophilin-2a (PKP2) interacts with Myh9 and Myh10.

PKP2 delocalization alters actomyosin cytoskeleton component organization. SARS-CoV-2 Nsp1 protein hijacks PKP2 from the desmosome into the soluble fraction where it is downregulated.

Viral Nsp1 collapses the actomyosin cytoskeleton and phenocopies the arrhythmogenic cardiomyopathy-related mutant R735X.

## Introduction

Coronavirus disease 2019 (COVID-19) is caused by the severe acute respiratory syndrome coronavirus 2 (SARS-CoV-2). In just a few months, COVID-19 has become a worldwide health problem of unprecedented spread since the 1918 Spanish flu. It is characterized by respiratory symptoms, but cardiac complications (Fried *et al*, 2020) including arrhythmias, heart failure, and viral myocarditis are also common, implicating myocardial injury as a possible pathogenic mechanism contributing to severe illness and mortality (Guo *et al*, 2020). Emerging evidence suggests that the SARS-CoV-2 infection compromises right ventricle (RV) function beyond inducing lung injury and acute respiratory distress syndrome. In fact, several studies show that RV dilation is strongly associated with in-hospital mortality, and that compromised RV function identifies higher risk patients with COVID-19 (Argulian *et al*, 2020; Li *et al*, 2020). Although the generalized inflammatory response (cytokine storm) caused by COVID-19 can induce a deleterious cardiac disfunction, the direct impact of SARS-CoV-2 infection on cardiac tissues is not well-understood. Like other coronaviruses, SARS-CoV-2 genome is made out of a singlestranded RNA, about ~30 kb in size, encoding for 26 proteins. The first open reading frames (orf) orf1a/b, located at the 5’ end, comprehend about two-thirds of the whole genome length, and encode polyproteins 1a and 1b (pp1a, pp1b). These polyproteins are processed into 16 non-structural proteins (NSPs) to form a replication-transcription complex that is involved in genome transcription and replication (Thi Nhu Thao *et al*, 2020). Amongst them is the N-terminal nonstructural protein 1 (Nsp1) that, similar to other coronaviruses, displays a analogous biological function suppressing host gene expression (Narayanan *et al*, 2008; Tohya *et al*, 2009). Nsp1 binds to the small ribosomal subunit and prevents canonical mRNA translation by blocking the mRNA entry tunnel (Thoms *et al*, 2020). Despite sharing over 84% of the amino acid sequence with the Nsp1 of the original SARS-CoV, the differences could hide new functions. Little is known about the role of Nsp1 in determined cell types and the consequences of recently described protein-protein interactions with host cell proteins like desmosomal *plakophilin-2 (PKP2)* (Gordon *et al*, 2020). Alterations affecting the cardiac isoform of *PKP2a*, which lacks the exon 6 (Gandjbakhch et al, 2011), account for 40–60% of genotype-positive patients with arrhythmogenic cardiomyopathy (ACM) (den Haan *et al*, 2009; Groeneweg *et al*, 2015; van Tintelen *et al*, 2006). Referred as a desmosomal disorder, ACM is a genetic disease of the heart muscle (Alcalde *et al*, 2014; Austin *et al*, 2019) that predisposes to sudden cardiac death (SCD) (Goff & Calkins, 2019), particularly in young patients and athletes (Coelho *et al*, 2019; Maron *et al*, 2004). For many years ACM has been known as arrhythmogenic right ventricular cardiomyopathy (ARVC) (Corrado *et al*, 2017a), showing autosomal-dominant inheritance and usually manifesting as impaired function of the RV (McKoy *et al*, 2000).

Desmosomes are dynamic intercellular junctions that maintain the structural integrity of skin and heart tissues by withstanding shear forces (Al-Jassar *et al*, 2013; Patel & Green, 2014). Actin is one of the major cytoskeletal proteins in eukaryotic cells and plays an essential role in several cellular processes, including mechano-resistance and contractile force generation (Koenderink & Paluch, 2018). Defective regulation of the organization of actin filaments in sarcomeres, owing to genetic mutations or deregulated expression of cytoskeletal proteins, is a hallmark of many heart and skeletal muscle disorders (Grimes *et al*, 2019).

Here, we uncover a new function of SARS-CoV-2 Nsp1 protein that drives inappropriate actomyosin cytoskeleton organization through a PKP2-dependent mechanism. By sequestering PKP2 from the desmosome, Nsp1 interferes with the turnover and localization of the actin-based motor proteins non-muscle myosin heavy chain II-A (Myh9) and B (Myh10). We identified these actomyosin components as key interactors of PKP2 cardiac isoform using biochemical and mass spectrometry approaches. The main role of Myh9 and Myh10 proteins is to orchestrate the mechanoenzymatic properties of stress fibers. Myosins execute numerous mechanical tasks in cells, including spatiotemporal organization of the actin cytoskeleton, adhesion, migration, cytokinesis, tissue remodeling, and membrane trafficking (Juanes-Garcia *et al*, 2016; Pecci *et al*, 2018; Vicente-Manzanares *et al*, 2009; Vicente-Manzanares *et al*, 2011; Vicente-Manzanares *et al*, 2007). Using atomic force microscopy together with fluorescence imaging, we show that expression of *PKP2* (c.2203C>T), encoding the R735X mutant found in ACM patients from different families (Alcalde *et al.*, 2014; Gerull *et al*, 2004) alters cell mechanical stiffness, and alike Nsp1, modifies actomyosin cytoskeleton. We found that *Nsp1* and *R735X* expression led to ultrastructural alterations in Myh9 and Myh10 and a rearrangement of the F-actin cytoskeleton, paired with reduced cell height and the collapse of the actomyosin cytoskeleton. These results suggest that PKP2 plays an important role in shape determination and actin homeostasis and that these functions can be modified both by the SARS-CoV-2 protein Nsp1 interaction and the ACM-related mutant R735X. All together our results suggest that Nsp1, through its interaction with PKP2 in the organization of actomyosin cytoskeleton, may induce a transient ACM-like phenotype in COVID-19 patients with cardiac infection.

## Results

### Mutant R735X expression alters actin cytoskeleton and cellular stiffness

We recently demonstrated that a C-terminal deletion *PKP2* mutant (*R735X*) operates as a gain-of-function protein in arrhythmogenic cardiomyopathy (ACM) (Cruz *et al*, 2015). To gain insight into the molecular mechanisms that induce the disease we studied the link between mutant PKP2 and cellular cytoskeleton. It is known that PKP2 loss impairs cortical actin remodeling and normal desmosome assembly (Godsel *et al*, 2010), which compromises cell mechanical properties like cellular stiffness (Puzzi *et al*, 2019). We measured cell stiffness in intact HL-1 cardiac cells by acquiring force-volume maps by atomic force microscopy (AFM). These force maps were used to represent spatial variation in Young’s modulus as a measure of local cell elasticity (stiffness) (Dufrene *et al*, 2017) in cell lines stably expressing PKP2 or mutant R735X (Figure 1A). The nanomechanical maps show that Young’s modulus was higher in regions with a high density of branched actin cytoskeleton networks and stress fibers. These observations were quantified by statistical analysis of Young’s modulus maps using a bottom-effect correction expression (Garcia & Garcia, 2018). The graphs show that the median value of the Young’s modulus measured in HL-1 PKP2 cells was higher than that measured in HL-1 R735X cells (Figure 1B). *Puzzi et al* using AFM in HL-1 cells showed that PKP2 knock-down was also associated with decreased cellular stiffness, as indicated by decreased Young’s modulus, when comparing to control groups (Puzzi *et al.*, 2019).These results support the observation that R735X cells lack most stress and F-actin fibers (Figure 1C). Images from the top central area of cells revealed a scarcity of F-actin filaments in the area above the nucleus of cells expressing the R735X mutant (46.05%±0.73% in PKP2 cells vs. 26.05%±1.48% in R735X cells; p<0.0001, ****; n=11) (Figure 1C). Together, these data show that the higher absolute levels of F-actin and stress-fiber formation in PKP2 HL-1 cell are associated with greater cell stiffness, whereas the lower cell stiffness in mutant R735X correlates with a reduction in F-actin and stress fiber disassembly.

**Figure 1.**
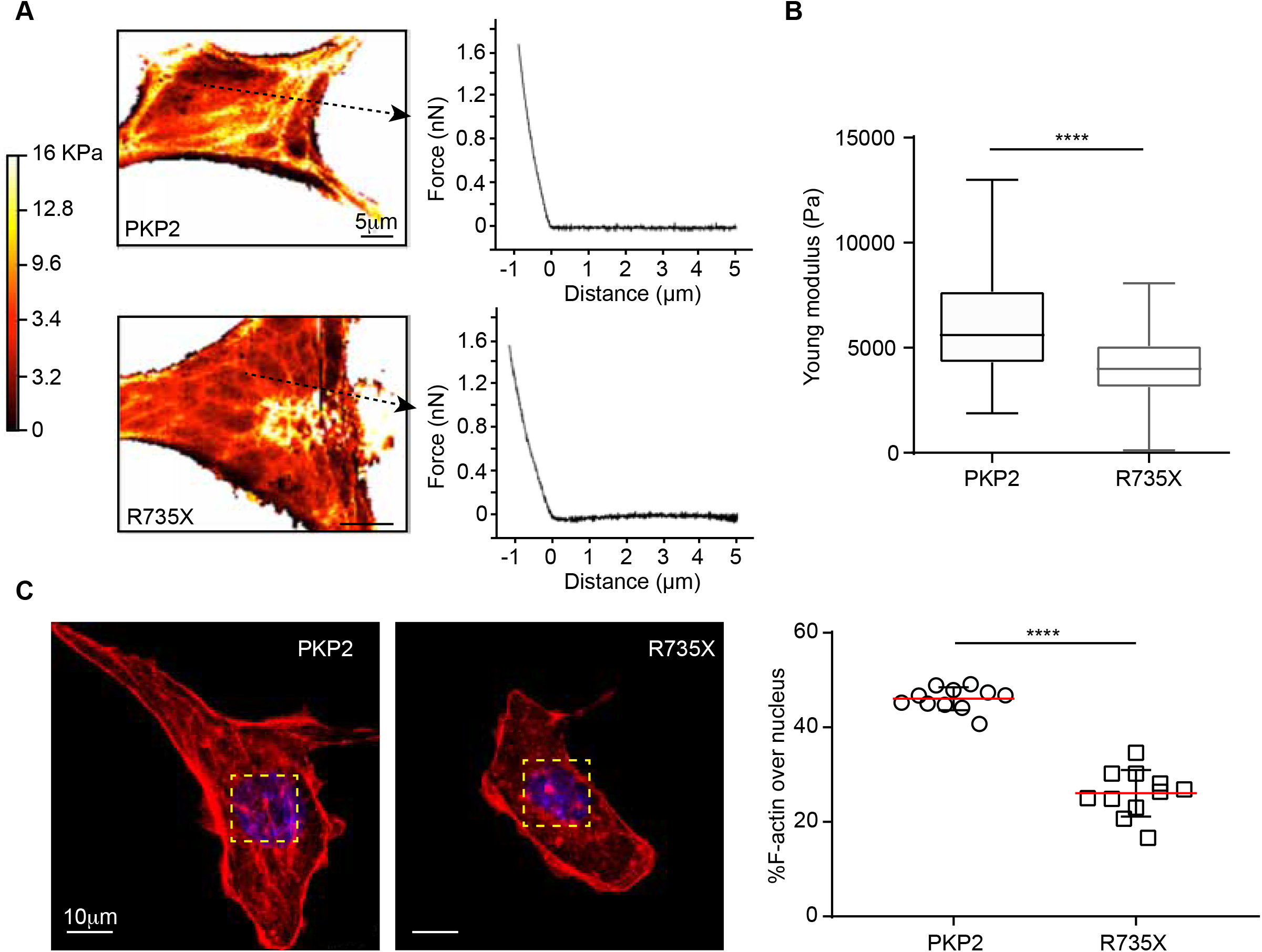
Mutant R735X alters actin organization and cellular stiffness. (A) Illustrative force maps of HL-1 cells expressing PKP2 or the R735X mutant and accompanying fitting force-distance curves, generated from atomic force microscope (AFM) measurements, showing the maximum indentation of the cell surface used to calculate Young’s modulus. (B) The box plot shows quantification of Young’s modulus as a measure of cell stiffness. Data are presented as median + minimum to maximum values (box limits); n=8080-93134, **** p<0.0001 (Mann-Whitney test). (C) Representative confocal images showing phalloidin-staining of the F-actin network (red) in HL-1 cells stably transfected with *pPB-PKP2* or *pPB-R735X*. Nuclei were stained with DAPI (blue). The yellow boxed areas demark the F-actin filaments overlying the nuclei in which phalloidin staining intensity was measured. Scale bars, 10 μm. The panel to the right shows quantitative analysis of F-actin abundance above the nuclei. Data are presented as mean ± SEM; n=11, **** p<0.0001 (unpaired Student *t* test).

### PKP2 interacts with the actomyosin proteins Myh9 and Myh10

To find candidates that could regulate actin cytoskeleton and cellular stiffness we explored the PKP2 interactome. We identified cardiac specific PKP2a (Gandjbakhch *et al.*, 2011), binding partners by pull-down of Halo-tagged PKP2 proteins followed by mass spectrometry (MS) analysis. PKP2-HaloTag pulldown products included previously identified interacting partners such as the desmosomal components desmocolin, plakoglobin, desmoglein, and desmoplakin together with novel interacting partners like myosin 9 (Myh9) and myosin 10 (Myh10), also known as non-muscle myosins NMIIA and NMIIB (Figure 2A). Interaction with Myh9 and Myh10 was validated by immunoprecipitation from cells expressing EGFP-PKP2 or EGFP-R735X (Figure 2B). These results support a direct link between desmosomal PKP2 and actomyosin cytoskeleton components.

**Figure 2.**
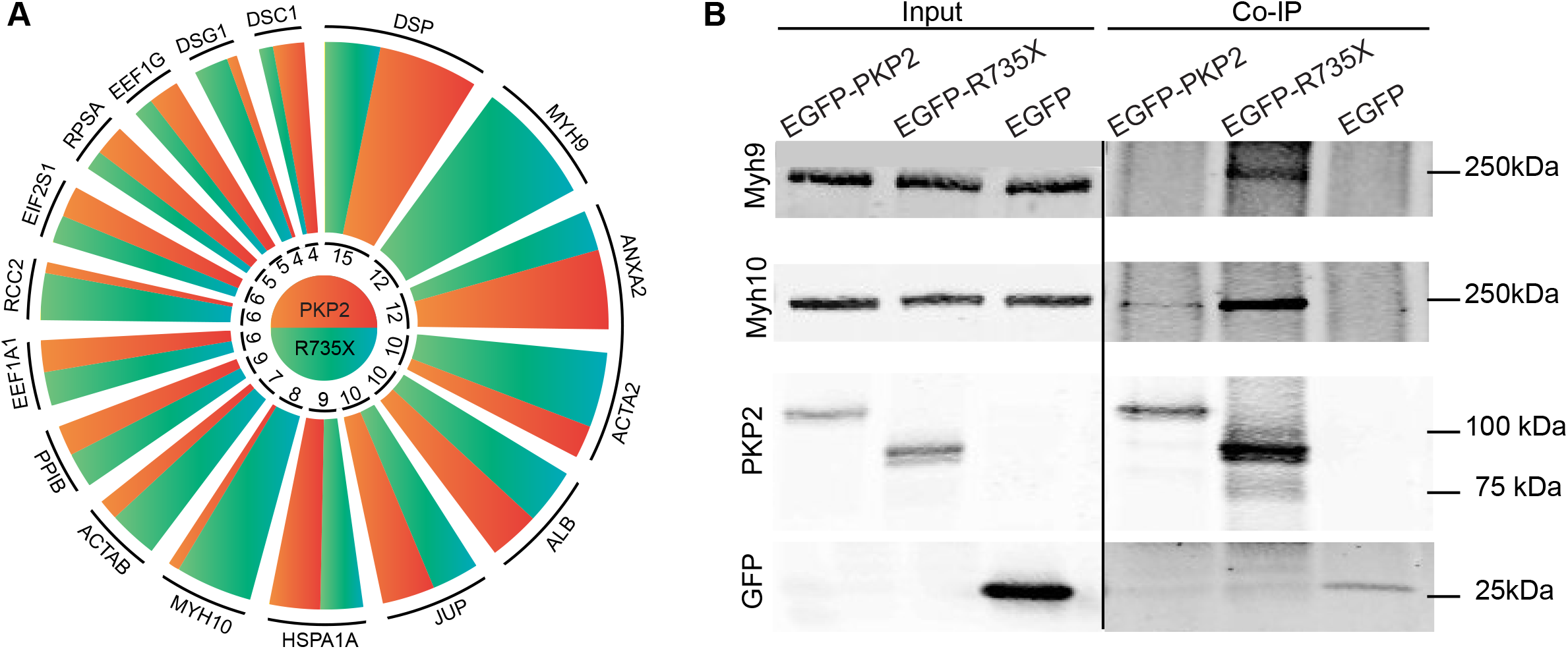
PKP2 interacts with Myh9 and Myh10. (A) Unique peptide counting of most abundant proteins identified in the HaloTag pull-down assays. Arc length is proportional to the total (PKP2 and R735X) number of unique peptides identified. (B) Western blot of proteins co-immunoprecipitated with EGFP, EGFP-tagged wild-type PKP2, or EGFP-tagged R735X mutant proteins prepared from HEK293T cells. Blots show PKP2 (wild-type and mutant versions), Myh9, Myh10, or EGFP. Blots from Input samples are also shown.

### PKP2 C-terminal deletion in mutant R735X alters its subcellular localization

Confocal microscopy revealed that most of the R735X mutant signal was located in the cytosol, displaced from the cell edge (Figure 3A). Image analysis in HeLa and HL-1 cells confirmed that R735X mutant was present at lower levels in the plasma membrane relative to the cytoplasm (Figure 3B). After transfection of cells with *pCAG-PKP2, pCAG-R735X*, or both plasmids, immunoblot analysis of cell fractions demonstrated that wild-type PKP2 associates with the plasma membrane, whereas the R735X mutant is predominantly detected in the cytoplasm and only occasionally at the membrane fraction (Figure 3C). These results demonstrate that the C-terminal region of PKP2 is important for proper protein localization at the border contact region. Also, highlight that PKP2 subcellular localization at the desmosome is not altered when co-expressed with mutant R735X PKP2 protein (Figure 3C). These data suggest that the gain-of-function mutant PKP2 in the cytosol has a new activity, as such function is not mediated by a deregulation of the levels or localization of wild-type PKP2.

**Figure 3.**
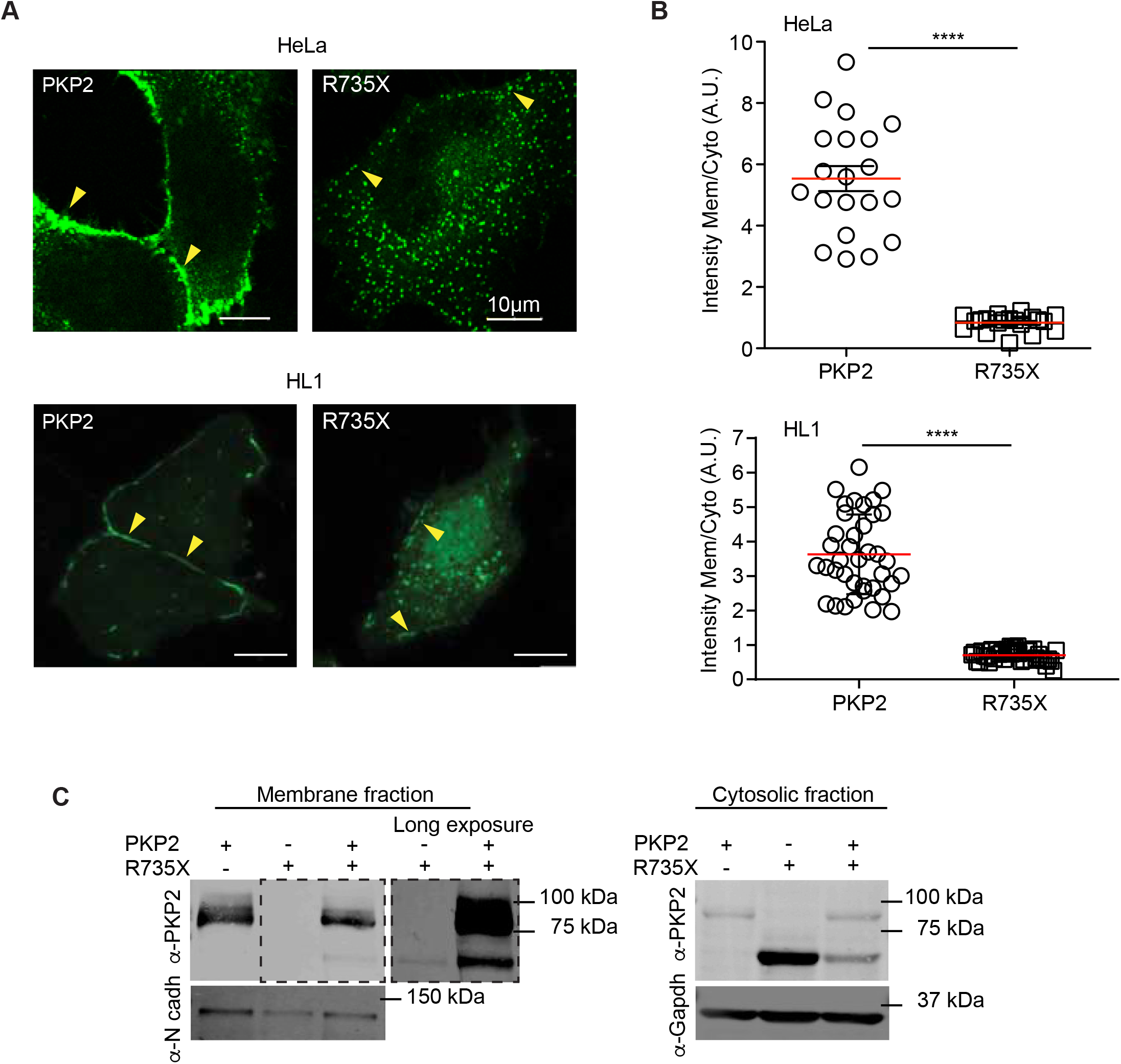
PKP2 C-terminal deletion alters its subcellular localization. (A) Subcellular localization of PKP2 (*pEGFP-PKP2*) and the R735X mutant (*pEGFP-R735X*). Arrowheads indicate enriched EGFP signal at the plasma membrane of HeLa (upper picture) and HL-1 cardiac cells (lower picture). Scale bars, 10 μm. (B) Graph shows the quantification of plasma membrane intensity levels normalized to cytoplasm intensity from (A). Data are presented as mean ± SEM; n=20-38, **** p<0.0001 (unpaired Student *t* test). (C) Immunoblot analysis of membrane and cytosolic fractions prepared from HEK293T cells. Symbols + and - indicate transfection or not with vectors encoding *PKP2, R735X*, or both variants. Blots show PKP2 (wild-type and mutant versions), N-cadherin (load control of membrane fraction) and GAPDH (load control of cytosolic fraction). Black dotted lines delimit short and long exposures of the same blot to detect R735X and PKP2 proteins.

### PKP2 localization defines Myh9 and Myh10 subcellular distribution

As Myh9 and Myh10 interact with mutant R735X in the cytoplasm, we hypothesized that PKP2 abnormal localization might disrupt actomyosin cytoskeleton organization. Firstly, we analyzed the precise subcellular distribution in cardiac HL-1 cells of endogenous actomyosin cytoskeleton (Myh9, Myh10 and actin) at cell periphery when using PKP2 or R735X EGFP N-terminal fusion protein (Figure 4A). Confocal imaging showed that PKP2 signal coincides with F-actin, and partially overlaps with Myh9 and Myh10. In fact, a representative fluorescence intensity profile (along a cross-section line from the plasma membrane to the cytoplasm; Figure 4A) of EGFP-tagged PKP2 (green line), together with Myh9 (red), Myh10 (blue) and F-actin (grey line), shows a structured disposition with a more prominent intracellular localization of Myh10, resulting in the absence of fluorescent colocalization with EGFP-PKP2 at the edge of the plasma membrane of the cell. Likewise, fluorescence of F-actin was found to largely colocalize with that of EGFP-PKP2. Line scan analysis shows that the vast majority of Myh9 and Myh10 filaments overlapped, except at the very leading edge where Myh9 is localized ahead of Myh10. By contrast, when R735X mutant is present, fluorescence of F-actin, Myh9 and Myh10 exhibited a full overlap, indicating inefficient distribution of these actomyosin cytoskeleton components at the cell edge (Figure 4B). To analyze the actomyosin elements distribution in the cell population we quantitatively evaluated the maximal signal intensity in multiple cells to confirm the effect of R735X (Figure 4C). It is important to notice that F-actin, Myh9 and Myh10 organization in R735X mutant differed from what can be observed in PKP2 shRNA cells. When PKP2 is absent only cortical actin is displaced from the periphery in agreement with previous reports (Godsel *et al.*, 2010) and Myh9 and Myh10 organization remain unaltered compared with control cells (Figure 4C). These results identify Myh9 and Myh10 as specific PKP2 interactors that provide a novel functional link between a desmosomal component localization and the actomyosin cytoskeleton organization.

**Figure 4.**
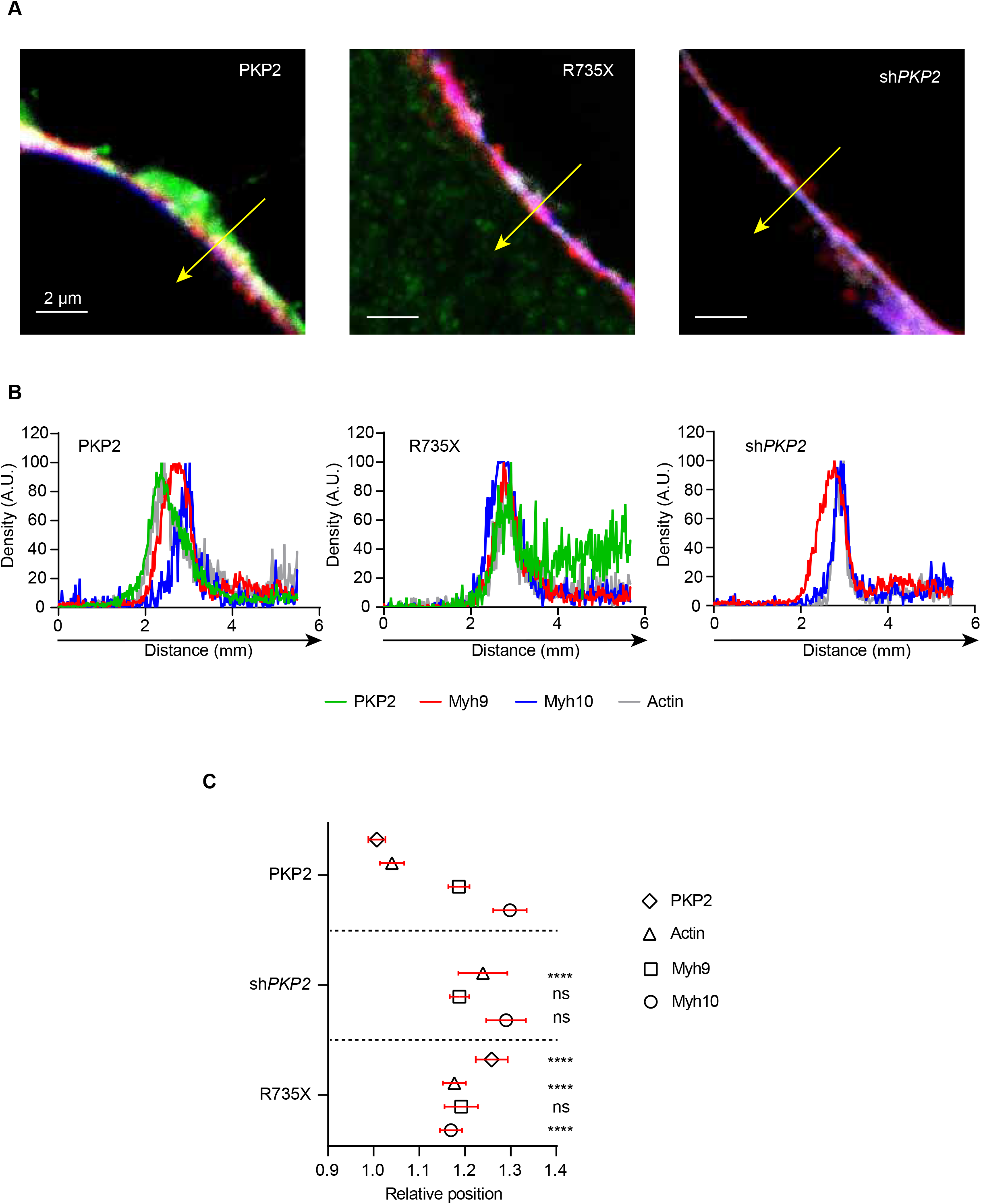
R735X expression modifies actomyosin component relative distribution. (A) Representative confocal images from immunofluorescence of PKP2 or R735X (green), Myh9 (red), Myh10 (blue) and Actin (grey) in HL-1 EGFP-PKP2, EGFP-R735X, and shRNA-*PKP2* stable cell lines. Scale bars, 2 μm. White arrows indicate the direction in which the intensity profiles of PKP2 (or PKP2 R735X mutant), Myh9, Myh10 and Actin on the membrane have been measured and the order in which the actomyosin components appear arranged (showed in C). (B) Representative fluorescence intensity profile graphs from (A) using the same colour code. (C) Box plot showing distance between the maximum intensity peaks of the intensity profiles between PKP2 or R735X, Myh9, Myh10 and Actin, in HL-1 cells stably expressing EGFP-PKP2, EGFP-R735X and sh*PKP2*. Data are presented as mean ± SEM; n=5, ns p>0.05, **** p<0.0001 (one-way ANOVA with the Tukey comparison post-test with PKP2 as the reference).

### Nsp1 sequesters PKP2 from the desmosome into the soluble fraction

A recent work from Krogan’s laboratory (Gordon et al., 2020) has also shown that the SARS-CoV2 nonstructural protein 1 (Nsp1) expressed in kidney HEK293 human cells as 2xStrep affinity tag fused protein pull downs PKP2 present in the lysates. Due to the preferential localization of PKP2 at the desmosome and the increased interaction of Nsp1-PKP2 in the soluble fraction (Gordon *et al.*, 2020) we speculated that Nsp1 hijacks PKP2 from the desmosome into the cytoplasm. To assess this hypothesis and to address whether Nsp1 delocalizes PKP2, we transfected human cells with EGFP-tagged *PKP2* and 2xStrep-tagged *Nsp1*. Representative confocal images of HeLa cell populations showed that under steady-state conditions EGFP-PKP2 signal was reduced and mobilized from the desmosome into the cytoplasm (Figure 5A). On the contrary, in the absence of viral Nsp1, wild-type transfected PKP2 localizes at the desmosomes around the cell perimeter. This effect of viral Nsp1 on PKP2 protein was observed in human and mouse cells (Figure 5B). Fluorescence intensity profiles spotted a lack of PKP2 fluorescent signal at the cellular cortex in cell cultures transfected with Nsp1 (Figure 5C). Consistently with the imaging analysis, we also confirmed that the expression of the viral protein Nsp1 decreases PKP2 absolute protein level and dramatically reduces the amount of protein in the insoluble fraction. When cell lysates were divided in NP-40 soluble and insoluble fractions we detected a significant decrease in the amount of PKP2 in the insoluble fraction (Figure 5D). These data suggest that PKP2 is not stably associated with the desmosomal complex in cells expressing Nsp1 and that PKP2 is unstable when displaced from the cellular cortex.

**Figure 5.**
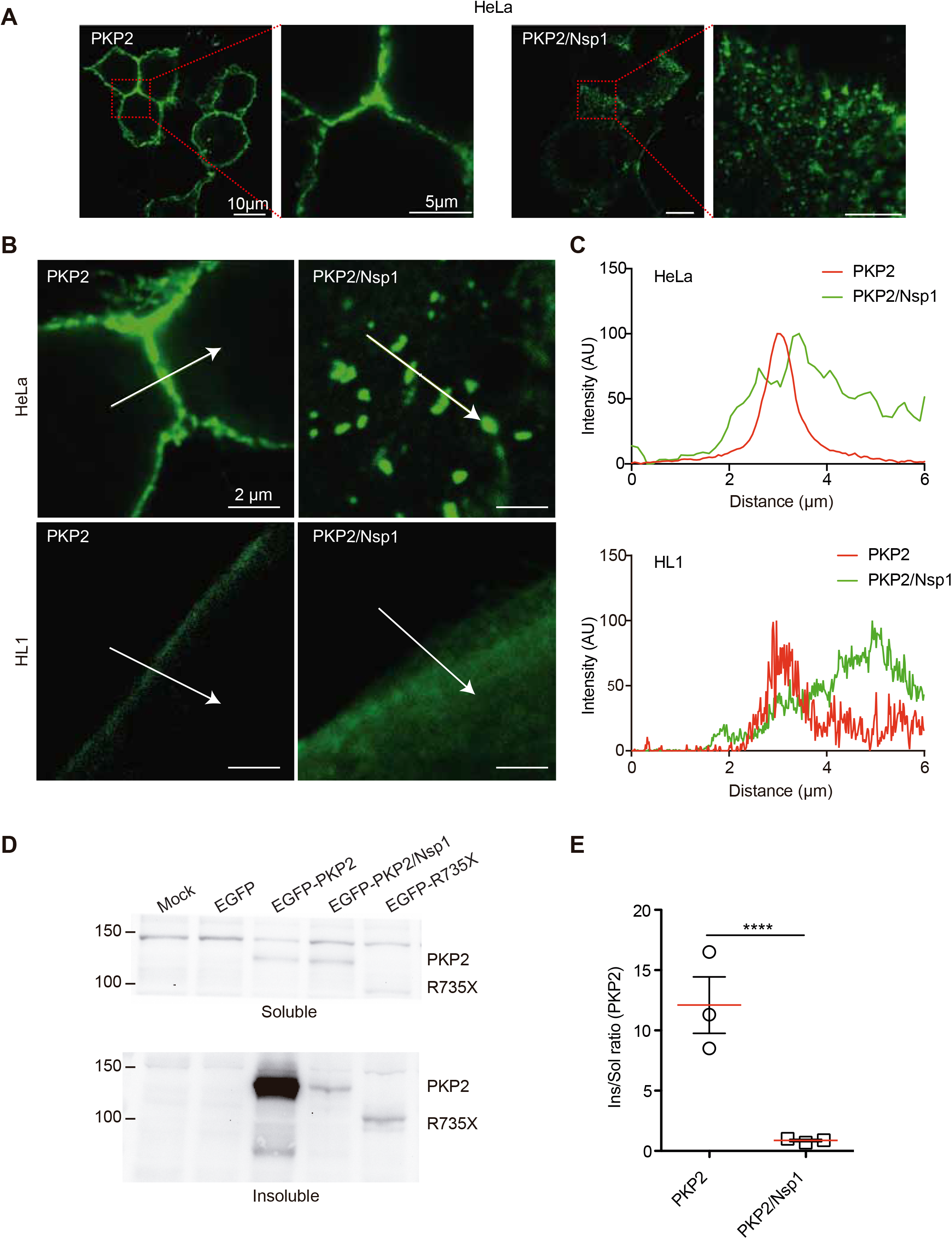
(A) Plasmids encoding *EGFP-PKP2* and *Nsp1* genes were transiently transfected in HeLa cells as indicated; 16 hours post-transfection EGFP-PKP2 cell signal was analyzed by confocal microscopy. Magnification of the indicated areas are shown on the right. Scale bars, 5 and 10μm respectively. (B) Subcellular localization of PKP2 (*pEGFP-PKP2*) with and without expression of viral Nsp1 protein. Arrowheads indicate the direction in which the intensity profiles of PKP2 have been measured in HeLa (upper picture) and HL-1 cardiac cells (lower picture). Scale bars, 2 μm. (C) Representative fluorescence intensity profile graph from (B). (D) Representative immunoblots showing the distribution of PKP2 from insoluble (Ins) and soluble (Sol) fractions of cells expressing PKP2 (*pEGFP-PKP2*) and Nsp1. (E) Graph shows PKP2 protein signal as an insoluble-soluble ratio calculated from densitometry analysis. Data are presented as mean ± SEM; n=4, **** p<0.0001 (unpaired Student *t* test).

### Nsp1 alike mutant R735X collapses the actomyosin cytoskeleton

We explored the effect of PKP2 and the R735X mutant when localized in the soluble fraction of the actin cytoskeleton. Firstly, we analyzed whether PKP2 displacement by Nsp1 phenocopies R735X mutant deregulation of Myh9 and Myh10 at the cellular edge (Figure 6A). Analysis of representative confocal images of EGFP-PKP2 HL-1 cells transduced Nsp1 lentivirus revealed that the subcellular distribution of endogenous actomyosin cytoskeleton (Myh9, Myh10 and actin) at cell periphery recapitulates the R735X organization (Figure 6B and 6C). Since changes in the actin cytoskeleton are intimately related to cell shape, well-organized actin in PKP2-expressing cells were shown by confocal microscopy z-stack imaging of fixed cells stained with phalloidin, whereas cells expressing the R735X mutant, or viral Nsp1 had a disorganized actin cytoskeleton (Figure 6D and 6E). Expression of either ACM-related R735X mutant or SARS-CoV-2 protein Nsp1 in the cells induce the collapse of the actomyosin cytoskeleton in mouse or human cells (Figure 6D and 6E). Transverse maximal projections of z-stack images in cells expressing PKP2 confirmed the organization of actin filament bundles into structures along the cell. In contrast, cells expressing the SARS-CoV-2 Nsp1 protein or R735X lacked this structure and the nucleus was close to the external plasma membrane with almost collapsed cytosolic space. We noted that both in Nsp1 expressing cells and R735X mutant, the actin distribution around the cell differed from that observed in PKP2 controls (Figure 6C) where actin fibers run over the nuclei. This difference in actin distribution was reflected independently of the cell type in the abnormal height of most R735X and PKP2-Nsp1 cells, which were shorter than control cells (PKP2, Nsp1 or mock transfected) (Figure 6D). We hypothesize that if the PKP2 delocalization effect on actin cytoskeleton is specific and based on an abnormal gain of function out of the desmosome, *PKP2* shRNA knockdown (sh*PKP2*), that do not induce PKP2 localization on the cytoplasm, and wild-type phenotypes should be the same. In fact, this is the case, as 70% *PKP2* knockdown does not alter cellular height in HL-1 cells (Figure 6E). All together, these results suggest that cytoplasmic localization of PKP2 either by a mutant involved in ACM development, or by Nsp1 hijack by SARS-CoV-2 infection can alter the integrity and assembly of the actin cytoskeleton by modulating the cortical distribution of Myh9 and Myh10.

**Figure 6.**
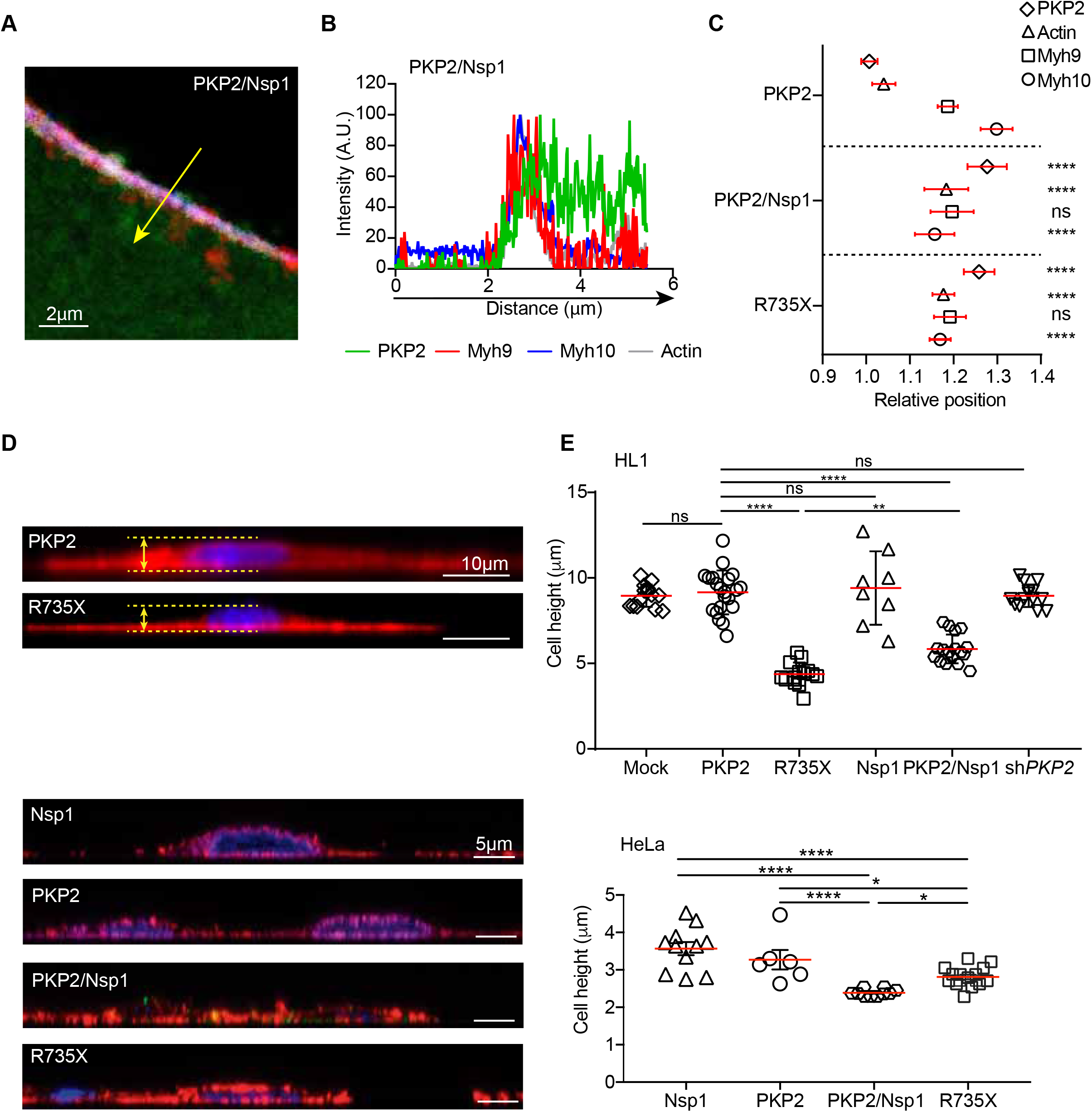
(A) Representative confocal image of PKP2 (green), Myh9 (red), Myh10 (blue) and Actin (grey) in HL-1 EGFP-PKP2, stable cell line infected with lentivirus encoding the SARS-CoV-2 protein Nsp1. Scale bars, 2 μm. Yellow arrow indicates the direction in which the intensity profiles have been measured and the order in which the actomyosin components appear arranged (showed in C). (B) Representative fluorescence intensity profile graph from (A) following the same colour code. (C) Box plot showing distance between the maximum intensity peaks of the intensity profiles between PKP2 or R735X, Myh9, Myh10 and Actin, in HL-1 cells stably expressing EGFP-PKP2, EGFP-R735X and expressing viral Nsp1 protein. Side views of HL-1 (D) and HeLa (E) cells stably expressing PKP2 or the R735X mutant generated by transverse maximal projection of phalloidin (red) and DAPI (blue) staining. The charts (right panel) plots maximum cell height corresponding to separation of the yellow lines in the HL1 cells images represented as an example. Data are presented as mean ± SEM; n=5-20, ns p>0.05, * p<0.05, ** p<0.001, and **** p<0.0001 (one-way ANOVA with the Tukey multiple comparison post-test).

## Discussion

Myocardial complications have been documented in a significant number of COVID-19 patients (Guo *et al.*, 2020; Shi *et al*, 2020), with a strong involvement of the right ventricle (RV) in individuals with poor prognosis. Our data suggest that the interaction between SARS-CoV-2 protein Nsp1 and desmosomal PKP2 in the soluble fraction of the cardiomyocytes could contribute to SARS-CoV-2-mediated cardiac damage and RV dysfunction by altering Myh9 and Myh10 homeostasis. At the molecular level we show that SARS-CoV-2 Nsp1 protein mimics the phenotype of the pathogenic mutation R735X, present in ACM families, by hijacking PKP2 and causing a collapse of the actomyosin cytoskeleton. Therefore, our results indicate that a direct cardiomyocyte invasion by the SARS-CoV-2 virus would induce a transient ACM-like phenotype, contributing to the RV deterioration observed in patients with COVID-19.

The most common subtype of ACM is arrhythmogenic right ventricular cardiomyopathy (ARVC), which is overrepresented in patients with mutations resulting in pre-mature termination of the PKP2 protein (Bhonsale et al, 2015; Lazzarini et al, 2015). Therefore, it is not surprising that pathogenic mutations in *PKP2* have been usually associated with the classical form of the disease that predominantly includes anomalous electrocardiograms and arrhythmias, with structural abnormalities that lead to a progressive global RV dysfunction (van Tintelen *et al.*, 2006). ACM presents very diverse penetrance and the same mutation may cause severe heart failure in one patient and no symptoms in another (Corrado *et al*, 2017b). These phenotypic differences can be explained due to environmental stressors like exercise (Cruz *et al.*, 2015; James *et al*, 2013) that define the final disease outcome. Under extreme exercise conditions the RV goes through a greater load increase compared with the left ventricle, where near-linear increase in pulmonary artery pressures predominantly contributes to a disproportionate increase in RV wall stress (La Gerche & Claessen, 2016). Given the profound effect of exercise on RV structure and function, it is reasonable that ACM develops in *PKP2* mutant carriers with compromised structural integrity of the myocardial cytoskeleton. Similarly, it is long recognized that RV dysfunction is frequently associated with moderate to severe acute respiratory distress syndrome (ARDS), which is one of the major determinants of COVID-19 mortality. Since most ARDS patients require mechanical ventilation, the uncoupling between the RV and pulmonary circulation under ventilation can also contribute to the fatal increase in RV stress. Our data demonstrate that SARS-CoV-2 Nsp1 protein alters PKP2 function by collapsing actomyosin cytoskeleton alike ACM-related mutant R735X. Given that the ACM development expressing R735X mutant is triggered by exercise (Cruz *et al.*, 2015), it is plausible the cardiac expression of Nsp1 contributes to aggravate RV cardiac dysfunction in patients with ARSD and/or mechanical ventilation. Thus, our data suggest that cardiac tissue infection by SARS-CoV-2 is overrepresented in COVID-19 patients with worst prognosis and higher death incidence. In fact, it has been documented that over consecutive COVID-19 autopsies, SARS-CoV-2 was found in the heart tissue in over 60% of the cases (Lindner *et al*, 2020).

Proteins do not exist in isolation, and the function of any given protein cannot be understood without considering its interactions with other proteins and its place in cellular interaction networks (Vidal *et al*, 2011). Mutations can affect interactions in many ways; they can completely wipe out all protein interactions; disrupt some interactions while retaining or strengthening others; or generate new interfaces. Loss of all interactions occurs most commonly through mutations that disrupt protein folding, leading to protein degradation (Sahni *et al*, 2015). However, this is not the case with the PKP2 R735X mutant, which can be detected (Figure 1), although does not accumulate as PKP2. It is possible that PKP2 once displaced from the desmosome, into the soluble fraction, is strongly regulated. In support of this hypothesis, we observed similar protein levels on the soluble fraction of mutant R735X and PKP2 dragged from insoluble structures by viral Nsp1 (Figure 4). Furthermore, both PKP2 and R735X proteins have a deleterious effect over actomyosin cytoskeleton organization what reinforces the relevance of regulating PKP2 levels outside the desmosome.

Our study reveals a desmosome-independent role of PKP2 in the regulation of Myh9, Myh10 and F-actin relative distribution at nanoscale level, exerted through protein-protein interaction. PKP2 gain-of-function mutant mainly localized in the cytoplasm alters actin remodeling required for accurate relative distribution and the maintenance of F-actin–derived biomechanical properties and cell architecture (Figures 4 and 6). This idea is reinforced by the fact that Nsp1 protein delocalizes PKP2 from the desmosome into the soluble fraction and deregulates the actomyosin network by downregulating and delocalizing Myh9 and Myh10.

Dominant inherited cardiomyopathy could be caused by loss-of-function (haploinsufficiency) or gain-of-function mechanisms. We identified a possible mechanism through which the *R735X* PKP2 variant could function as a gain-of-function mutation and not a simple loss-of-function allele. We hypothesize that the pathogenic R735X variant functions by interrupting or rewiring highly connected interaction networks to disturb F-actin homeostasis. We predict that gain of function will also be the mode of action of other C-terminally truncated PKP2 variants stable to retain Myh9 and Myh10 interaction and to disrupt cell-to-cell subcellular localization. The ClinVar database (https://www.ncbi.nlm.nih.gov/clinvar/), part of the NCBI Entrez system, attempts to establish relationships between gene variants and phenotype. Among them, 34 are nonsense mutations and 23 span different portions of the 8 armadillo domains in PKP2. Remarkably, all nonsense *PKP2* variants are classified as pathogenic or likely pathogenic, underlining the functional importance of the PKP2 C-terminal domain, although the mechanism by which they induce pathogenicity would be diverse. Taking into context our results when Nsp1 is present, and the effect of PKP2 delocalization, it is possible that all PKP2 mutants that lack the ability to stably localize in the desmosome will share similarities in their phenotypes when they maintain their interaction with Myh9 and Myh10.

The phenotype associated with the R735X mutant is largely different from wild-type PKP2 localized at the cell edge, revealing a major role for relocalization into the soluble fraction of the mutant variant in actomyosin homeostasis. This idea is strengthened by the fact that delocalization of PKP2 into solution by viral Nsp1 alters actomyosin components to collapse the cellular cytoskeleton. Therefore, the delocalization of PKP2 and the disorganization of Myh9 and Myh10 at the cellular cortex could explain the aberrant organization of the cytoskeleton observed in cells expressing Nsp1 or the ACM-related mutant R735X. In summary, our data shows that PKP2 plays a central role in cytoskeleton homeostasis through the interaction and localization of actomyosin components Myh9 and Myh10, and that function is disturbed by the SARS-CoV-2 protein Nsp1 with similar consequences to ACM pathogenic mutations observed in PKP2.

## Materials and methods

### Cell lines

The HEK293T (ATCC, CRL-3216) and HeLa (ATCC, CCL-2) cell lines were maintained in DMEM (GIBCO) supplemented with 10% FBS, 1% penicillin/streptomycin, and 2 mM L-glutamine. The atrial cardiomyocyte cell line HL-1 (Sigma, Aldrich) was maintained in Claycomb medium (Sigma Aldrich) supplemented with 10% FBS, 1% penicillin/streptomycin, and 2 mM L-glutamine, as previously described (Claycomb *et al*, 1998). HL-1 cells were seeded on plates coated with 0.02% gelatin/fibronectin (Sigma Aldrich). Cell lines were maintained at 37°C with 5% of CO_2_.

### Transfection and stable cell lines

Transient transfection with cDNAs encoding EGFP, EGFP-tagged PKP2 and R735X, Nsp1 proteins were performed in 35-mm tissue culture dishes (MatTek) using the jetPRIME® reagent according to the manufacturer’s protocol (Polyplus transfection®). Stable HL-1 cell lines were generated using the *PiggyBac* transposon system. Cells were transfected with plasmids expressing *PKP2*, *shPKP2, EGFP-PKP2, R735X, EGFP-R735X* or *R735X-EGFP* together with the *pPB-transposase* (Cadinanos & Bradley, 2007). Cells were then selected for geneticin resistance (G418, ThermoFisher Scientific) and expanded for further experiments.

### Atomic Force Microscopy (AFM)

The AFM experiments were performed with a commercial instrument (JPK NanoWizard 3, JPK Instruments AG, Berlin, Germany) mounted on an Axio Vert A1 inverted microscope (Carl Zeiss, Oberkochen, Germany). For these experiments, HL-1 cells stable expressing PKP2 or R735X were maintained in Claycomb cell culture medium. To ensure the cells stayed alive and adherent during the force spectroscopy experiments, measurements were made at a constant temperature of 37°C. Cell studies were conducted using specially adapted Biotool Cell XXL cantilevers (NanoandMore, Germany) with a nominal spring constant of 0.1 N/m and a height of 15 μm; the use of extra-long tips of ~15 μm minimizes accidental contact of the cell with the cantilever beam body. The half-cone angle was ~12°, and the nominal radius at the tip apex was 25 nm. The upper cantilever surface was gold-coated to improve the signal-to-noise ratio in the deflection signal. To control the force applied on the cell, the deflection sensitivity was calibrated on a Petri dish. The spring constant (0.09-21.1 N/m) was calculated using the thermal noise method (Lozano *et al*, 2010). Force-volume maps (Dufrene *et al.*, 2017) were generated for whole cells by acquiring force-distance curves (128×128 pixels^2^) over a 30 μm^2^ area. The tip sample distance was modulated by applying a triangular waveform. Each individual force-distance curve was acquired at a velocity of 100μm/s (8Hz) and a range of ~6 μm. To prevent sample damage, the maximum force applied to cells was 2 nN.

Maps were analyzed with in-house software written in Python. We analyzed 8 HL-1 cells expressing PKP2 and 5 cells each expressing the R735X and R735X-EGFP variants. The program includes bottom-effect corrections for a conical tip to correct for finite cell thickness (Garcia & Garcia, 2018).

### HaloTag pull-down

PKP2 isoforms were expressed in HEK293T cells as N-terminal Halo tag fusion proteins. Cell pellets were lysed using Mammalian Lysis Buffer (G9381, Promega). The bait-prey complexes, containing the PKP2–Halo-tagged fusion protein (bait) and the potential binding partners (prey), were pulled down using HaloLink resin (Promega Madison, WI) and extensively washed in buffer containing 100 mM Tris (pH 7.6), 150 mM NaCl, 1 mg/ml BSA, and 0.05% IGEPAL® CA-630 (octylphenoxypolyethoxyethanol, I3021, Sigma-Aldrich, Oakville, ON). Purified bait-prey protein complexes were digested overnight with TEV protease at 4°C to release halo-linked PKP2 protein, and the tag-free protein complexes were isolated with a His-Trap-Spin column.

### Protein digestion, mass spectrometry and peptide identification

The eluted protein complexes were in-gel digested with trypsin as described previously (Bonzon-Kulichenko *et al*, 2011), and the resulting peptides were analyzed by liquid chromatography coupled to tandem mass spectrometry (LC-MS/MS), using an Easy nLC-1000 nano-HPLC apparatus (Thermo Scientific, San Jose, CA, USA) coupled to a hybrid quadrupole-Orbitrap mass spectrometer (Q Exactive HF, Thermo Scientific). The dried peptides were taken up in 0.1% (v/v) formic acid and then loaded onto a PepMap100 C18 LC pre-column (75 μm I.D., 2 cm, Thermo Scientific) and eluted on line onto an analytical NanoViper PepMap 100 C18 LC column (75 μm I.D., 50 cm, Thermo Scientific) with a continuous gradient consisting of 10–35% B (80% acetonitrile, 0.1% formic acid) for 60 min at 200 nL/min. Peptides were ionized using a Picotip emitter nanospray needle (New Objective). Each mass spectrometry (MS) run consisted of enhanced FT-resolution spectra (120,000 resolution) in the 400–1500 m/z range followed by data-dependent MS/MS spectra of the 20 most intense parent ions acquired during the chromatographic run. For the survey scan, the AGC target value in the Orbitrap was set to 1,000,000. Fragmentation in the linear ion trap was performed at 27% normalized collision energy, with a target value of 100,000 ions. The full target was set to 30,000, with 1 microscan and 50 ms injection time, and the dynamic exclusion was set to 0.5 min. The MS/MS spectra were searched with the Sequest algorithm in Proteome Discoverer 1.4 (Thermo Scientific). The UniProt human protein database (March 2017, 158,382 entries) was searched with the following parameters: trypsin digestion with 2 maximum missed cleavage sites; precursor and fragment mass tolerances of 800 ppm and 0.02 Da, respectively; Cys carbamidomethylation as a static modification; and Met oxidation as a dynamic modification. The results were analyzed using the probability ratio method (Martinez-Bartolome *et al*, 2008), and a false discovery rate (FDR) for peptide identification was calculated based on search results against a decoy database using the refined method (Navarro & Vazquez, 2009).

### Lentivirus production

Nsp1-expressing lentiviral particles were produced by cotransfection of *psPax.2* (gag-pol), *pMD2-G* (env) and *pLVX-EF1a-Nsp1-2xStrep-IRES-Puro* plasmids into HEK293T cells with calcium phosphate. Viruses were harvested 72 h posttransfection, filtered through 0.45 um PES filter and concentrated by ultracentrifugation prior to titration by qPCR.

### Immunostaining and imaging analysis

HeLa and HL-1 cell lines were fixed with 4% formaldehyde for 10 minutes, then permeabilized for 90 minutes at room temperature with 0.1% Triton X-100, and finally stained with Phalloidin-iFluor 594 Reagent (ab176757, Abcam). Cells were washed with phosphate buffered saline before addition of DAPI (ThermoFisher Scientific) and were examined under a Leica confocal laser scanning microscope. Z-stack images were captured and prepared as maximal projections. Cell height was measured from maximal projections of transverse Z-stack images, taking account the size (in μm) of the Z-stack. The percentage of actin filaments overlying the nucleus was calculated from the phalloidin intensity measured with Fiji software in all Z-stack images above the nucleus.

For actomyosin components distribution analysis HL-1 EGFP stable cell lines were seeded at a density of 1×10^4^/cm^2^ and cultured for 24 h. After 15 min fixation with 4% PFA and permeated with 0.5% Triton X-100 PBS, cells were blocked with 10% normal goat serum (NGS) in 0.1% Triton X-100 PBS for 1 hour at room temperature. Then, cells were incubated with chicken α-GFP antibody (1:200, GFP-1010, Aves Labs), rabbit α-Myh9 antibody (1:200, GTX113236, GeneTex) and mouse α-Myh10 antibody (1:200, GTX634160, GeneTex) in 0.1% Triton X-100 PBS containing 10% NGS overnight at 4°C. After washing with 0.1% Triton X-100 PBS, cells were incubated with secondary antibodies for 1 hour at room temperature: Alexa fluor 488 IgG goat antichicken (1:500, A32931, Invitrogen), Alexa fluor 647 IgG goat anti-rabbit (1:500, A32733, Invitrogen) and Alexa fluor 405 IgG goat anti-mouse (1:500, no. A31553, Invitrogen). After washes with 0.1% Triton X-100 PBS, actin was stained with Phalloidin-iFluor 594 Reagent (ab176757, Abcam) for 90 minutes at room temperature. After washes in 0.1% Triton X-100 PBS, samples were mounted in Mowiol mounting medium (Mowiol 4-88, Glycerol, 200 mM Tris-HCl pH 8.5 and 2.5% 1,4-diazabicyclo-[2,2,2]-octane). Fluorescence images were obtained with a Leica SP8 confocal microscope. To study the distribution of PKP2 (or R735X), Myh9, Myh10, and Actin proteins, a 6 μm line over the plasma membrane was drawn from extracellular space to cytoplasm and profiles of intensity for all of them were analysed. Peaks of intensity for these four proteins were identified and the distance between PKP2 and Myh9, Myh10 or Actin, were measured and normalized to PKP2.

To calculate ratio membrane-cytoplasm fluorescence z-stacks images from HL-1 cells expressing EGFP-PKP2, EGFP-R735X or R735X-EGFP were acquired with a Leica SP8 confocal microscope with HC PL APO 100x/1.4 oil objective. Regions of interest (ROIs) were drawn over maxima projections images to define the plasma membrane and the cytoplasm, excluding the nucleus and/or vacuoles. The ratio plasma membrane-cytoplasm intensity was calculated to normalize the intensity of the plasma membrane to the level of expression on each single cell.

### Cell fractionation

After 24 hours, transfected cells were washed once with ice-cold PBS and after dislodged by scraping using Buffer NP40 (50 mM Tris-HCl pH7,5; 150 mM NaCl; 1% Nonidet P 40 substitute). Cells were lysed for 30 min at 4°C on a rotator and the lysates were cleared by centrifugation (13000 rpm for 30 min at 4°C). Pellet was resuspended in Buffer NP40 supplemented with 1% Triton X-100 and 1X protein loading Buffer (40 mM Tris-HCl pH 7,5, 2% SDS, 2 mM β-mercaptoethanol, 4% glycerol, 0,02% Bromophenol Blue).

For plasma membrane protein extraction cells were washed once with ice-cold PBS and dislodged by scraping. Plasma membrane proteins were extracted using the Plasma Membrane Protein Extraction Kit from Abcam. Proteins were separated from membrane and cytoplasmic extracts on 8% SDS-PAGE gels and then western blotted with antibodies against PKP2 (EB10841, Everest Biotech), N-cadherin (sc-59987, Santa Cruz Biotechnology), and GAPDH (sc-32233, Santa Cruz Biotechnology). Secondary antibodies were anti-goat (ABIN2169607, Antibodies online) and anti-mouse (ABIN6699027, Antibodies online) as appropriate. Immunoblots were developed with the Odyssey Imaging System.

### Co-Immunoprecipitation

HEK293T cells were transfected with *pEGFP-N1, pEGFP-PKP2*, or *pEGFP-R735X*. After 48 hours, cells were washed with ice-cold PBS and dislodged by scraping. Proteins were extracted in NP-40 buffer (150 mM NaCl, 50 mM Tris-HCl pH 8.0, 1% NP-40, and protease and phosphatase inhibitors). Samples were incubated with shaking for 2 hours at 4°C and then centrifuged for 30 minutes at 20,000g at 4°C. All protein samples were quantified by the Lowry method (BioRad), and 1mg and 20μg of each sample were used for coimmunoprecipitation (Co-IP) and input, respectively. Co-IP was performed with Dynabeads® A (10008D, ThermoFisher Scientific). For each sample, 50 μl Dynabeads were washed four times with NP-40 buffer. After this, rat IgG1 anti-EGFP (kindly provided by the Monoclonal Antibody facility at the CNIO, Spain) were added to the Dynabeads at 1:100 dilution and incubated with shaking for 45 minutes at 4°C. The antibody was then removed, and 1mg of protein per sample was added to the Dynabeads, followed by incubation overnight with shaking at 4°C. Unbound proteins were removed by washing the Dynabeads four times in NP-40 buffer. Bound proteins were eluted with 30 μl loading buffer (10% SDS, 10 mM β-mercapto-ethanol, 20% glycerol, 200 mM Tris-HCl pH 6.8, 0.05% Bromophenolblue), heated for 15 minutes at 95°C, separated on 8% SDS-PAGE gels, and western blotted with antibodies against MYH9 (GTX633960, GeneTex), MYH10 (3404S, Cell Signaling Technology), and EGFP (632380, Clontech). Secondary antibodies were anti-mouse (ABIN6699027, Antibodies online) and anti-rabbit (ABIN5563398, Antibodies online), as appropriate. Immunoblots were developed with the Odyssey Imaging System.

### Statistics

No data were excluded from the analysis. All data were analyzed by one-way ANOVA with the Tukey multiple comparison post-test, two-way ANOVA, unpaired Student *t* test, or Mann-Whitney test. Error bars represent SEM. Statistical significance of differences was assigned as follows: * p<0.05, ** p<0.01, *** p<0.001, **** p<0.0001, and ns p>0.05.

## Author Contributions

C. M-L., M.R-M., S.S., N.G-Q. and J.A.B. designed and interpreted the experimental work to characterized the PKP2 mutant R735X models, and the Nsp1 cells with the help of A.G-G., M.L., and MI-G.. C.M-L., D.M-P., C.S-R. D. S-R., and A.G-G. constructed the plasmids and generated the cell lines. E.C. was responsible for mass spectrometry analyses with the help of M.R-M. D.S. performed the AFM experiment and analyzed the data. R.G. designed, analyzed, and interpreted the AFM experiment. J.A.B. conceived the project, and wrote the manuscript. All authors discussed the results and commented on the manuscript.

